# Functionally-structured Bayesian model for localizing neural activity and information in magnetoencephalography signals

**DOI:** 10.1101/2025.07.02.662527

**Authors:** Kai Miyazaki, Shun Nirasawa, Naoki Ishibashi, Kazuaki Akamatsu, Okito Yamashita, Yoichi Miyawaki

## Abstract

Magnetoencephalography (MEG) is a noninvasive method that can measure human brain activity with high temporal resolution. However, the spatial resolution of MEG is not sufficient to reveal neural mechanisms. Although MEG source estimation overcomes this problem to some extent, the combination of MEG source estimation and multivariate analysis results in “information spreading”, where significant predictions are observed in brain areas outside the true signal source location. This paper describes a model that achieves high source estimation accuracy and suppresses information spreading simultaneously. The proposed approach is based on a Bayesian estimation model that utilizes functional structure of the human brain. We compare the performance of the proposed model with simulated data generated under various signal-to-noise ratio conditions. The results show that the functionally-structured Bayesian model achieves source estimation accuracy that is better than that of conventional source estimation models. Additionally, the comparison of information spreading among the models reveals that our model outperforms the conventional ones. These results suggest that information spreading in the MEG source estimation can be suppressed while maintaining high source estimation accuracy.

## 1. Introduction

The human brain encodes information with neural activity patterns that change at fine spatial and temporal scales. Neuroimaging techniques thus require a sufficient spatiotemporal resolution for tracking such neural activity patterns. The spatial resolution of functional magnetic resonance imaging (fMRI) is high enough to discern the cortical layer structure at a submillimeter level. However, its temporal resolution is poor because its signal depends on the blood-oxygen-level change. On the other hand, magnetoencephalography (MEG) measures the magnetic field generated by electrical neural activity inside the brain using magnetic sensors at a high temporal resolution, but the spatial resolution of MEG is insufficient because the magnetic sensors are placed above the scalp and each sensor measures the magnetic field originating from multiple brain areas. At present, therefore, there are no neuroimaging methods that can track neural activity patterns at a sufficient spatiotemporal resolution.

To resolve this issue, previous studies have attempted to estimate the source locations in the human brain that correspond to measured MEG signals (Hämäläinen, M S and Ilmoniemi, R J, 1994; Iwaki and Ueno, 1998; Sato et al., 2004). Various models based on this MEG source estimation approach have been developed and tested. The most widely used MEG source estimation approach involves relaxing the ill-posedness, which is caused by the number of sensors being much smaller than the number of candidate source locations in the brain, by using a regularization technique. Minimum norm estimation (MNE) (Hämäläinen, M S and Ilmoniemi, R J, 1994) uses squared regularization, yielding a stable but spatially spread solution. In contrast, minimum current estimation (MCE) (Tibshirani, 1996) and automatic relevance determination (ARD) (Bishop, 2006; MacKay, 1992) apply regularization that promotes sparse solutions. The high-spatial-resolution fMRI information can also be used to improve the source estimation accuracy. For example, activity information such as the blood-oxygenation-level-dependent (BOLD) signal contrast of fMRI data can be used as prior information for the MEG source estimation. Variational Bayesian multimodal encephalography (VBMEG) (Sato et al., 2004) uses the activity information as a hierarchical prior within a Bayesian estimation framework. Although this model has achieved accurate source estimation to some extent, it remains unclear how accurately the estimated cortical current restores information represented in the original neural activity patterns.

To examine whether MEG source estimation can restore information represented by neural activity patterns, one reasonable approach is to try to predict the original information from the estimated cortical current. This approach is known as “decoding” and have been applied in many neuroimaging data (Cox and Savoy, 2003; Miyawaki et al., 2008). Sato et al. (2018) combined MEG source estimation with decoding and showed that the original information can be accurately predicted from the estimated cortical current. However, they also found statistically significant predictions even outside the originally active source areas. In other words, information representation was unexpectedly detected even in the brain areas that do not contain any informative activation. This phenomenon is called “information spreading,” easily leading to false-positive interpretations when combining MEG source estimation with decoding analysis to identify the brain areas that represent the target information. Sato et al. (2018) evaluated various MEG source estimation models in terms of source estimation accuracy and the degree of information spreading. The results showed that source estimation models that tend to produce spatially spread estimation like MNE and beamformer yielded information spread in the wide brain areas beyond the original active source areas. Furthermore, even VBMEG that revealed the most accurate source estimation tightly localized in the active source area showed information spreading in the wide brain areas as MNE and beamformed did. In contrast, MCE that promotes sparse source estimation did not show wide information spreading as other models did, but this is probably because the source estimation itself failed even in the original active source area as well as the outside areas (the source estimation accuracy was worst in the tested models). Therefore, at present, there are no MEG source estimation models that can achieve both high source estimation accuracy and the suppression of information spreading.

To address this issue, we focus on the fact that the human brain structure is subdivided into the multiple functionally distinct groups, and develops a model that incorporates the functional structure information into the MEG source estimation model. Such functional structure is defined in various human brain atlases (Destrieux et al., 2010; Glasser et al., 2016; Yeo et al., 2011, and more), showing the groups of the cortical locations that are supposed to have similar functions achieved by similar neural activity. In contrast, conventional source estimation models typically treat each cortical location independently and do not consider the similarity of neural activity among the cortical locations within the same functional area. We hypothesize that incorporating this functional structure information into MEG source estimation may improve the source estimation accuracy and suppress information spreading beyond the target source area. Then, we group cortical points based on the functional structure information and construct a model that can be updated by a group-by-group fashion. This model is expected to better capture the tendency for neighboring cortical points in the same functional area to exhibit common activity patterns. This would lead to lower correlations in estimated current between different cortical areas and consequently reduce the information spreading beyond the target area. The grouped least absolute shrinkage and selection operator (gLasso)(Lim et al., 2017; Yuan and Lin, 2006) uses the functional structure information to define parameter groups for regularization, promoting sparse solutions at the group level. Although this model has improved performance over MNE, it has not been compared with Bayesian approaches using the same functional information. A similar approach is grouped automatic relevance determination (gARD) (Yu et al., 2015), which incorporates group information in a different way as a prior within a Bayesian framework. gARD grouped EEG signals from the same channel to predict the presence or absence of the P300 component from the spatiotemporal patterns of EEG signals. However, it remains unclear whether gARD framework is useful for MEG source estimation, especially when incorporating physiologically-relevant group information according to the functional organization of the human brain. In this study, we propose a functionally-structured Bayesian approach and compare the performance between these frameworks. Through this approach, we aim to achieve more accurate source estimation accuracy and better control of information spreading simultaneously.

## 2. Method

To evaluate the performance of source estimation, we tested multiple models, including the functional structure model, using both simulation and real data as follows.

### 2.1. Simulation

#### 2.1.1. Participant

One individual participated in the magnetic resonance imaging (MRI) experiment to acquire an anatomical image of the participant’s brain, which were used to extract the cortical surface model. The participant gave written informed consent before the experiment. The procedure was approved by the institutional review board of the University of Electro-Communications, Advanced Telecommunications Research Institute International (ATR) Brain Activity Imaging Center, and Ethical Review Board of Osaka University Hospital. The experiment was conducted under corrected vision with MRI-compatible glasses.

#### 2.1.2. MRI experiment

Anatomical images were obtained using an MRI system (3-Tesla MAGNETOM Prismafit, Siemens) located at the ATR Brain Activity Imaging Center. The magnetization-prepared rapid-acquisition gradient-echo (MP-RAGE) sequence (208 sagittal slices; TR, 2,250 ms; TE, 3.06 ms; FA, 9 deg; FOV, 256 × 256 mm; voxel size, 1.0 × 1.0 × 1.0 mm) to was used to acquire the anatomical images.

#### 2.1.3. Head position measurement and registration

First, a 3D scanner and position sensor system (Polhemus Cobra) were used to measure the shape of the participant’s head and the positions of the five marker coils (three on the forehead and two on each earlobe) that were used to calibrate the head position relative to the MEG sensors inside the MEG scanner (PQ1160C, Yokogawa Electric Co., Japan). Then, the participant was asked to lie on their back in the MEG scanner and remain as still as possible. The current was then applied to the five marker coils to generate a magnetic field, and the magnetic field was measured with 160-channel superconducting quantum interference device (SQUID) sensors in the MEG scanner. The positional relationship between the MEG sensors and the marker coils was obtained by solving an inverse problem from the measured magnetic field. Using the position information of the marker coils, the participant head location relative to the MEG sensor locations was calculated. Finally, the high-resolution anatomical MRI image of the participant was aligned to the participant head locations in the MEG scanner.

#### 2.1.4. Calculation of MEG signals

The MEG signals were calculated by a forward model derived from the positional relationship between the signal source and the MEG sensors. The forward model can be expressed as

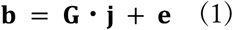

where 𝐛 (ℝ^𝑁×1^ vector; 𝑁, the number of MEG sensors) is the measured MEG signal, 𝐆 (ℝ^𝑁×M^ vector; 𝑀, the number of vertices on the cortical surface) is a leadfield matrix that represents how large MEG signals are produced at each sensor location by each cortical current, 𝐣 (ℝ^𝑀×1^ vector) is the cortical current, and 𝐞 (ℝ*^N^*^×1^ vector) is the sensor noise. 𝐆 was calculated using the VBMEG software. Procedures to set 𝐣 and 𝐞 is described in the next subsection.

#### 2.1.5. Regions of interest

In this study, we used a standard brain model provided by the Human Connectome Project (HCP) dataset (Glasser et al., 2016), consisting of 180 functional regions in the cortical surface of each hemisphere (360 in the both hemispheres) as regions of interest (ROIs). To assign these ROIs for the participant cortical surface, we used the Freesurfer software (http://surfer.nmr.mgh.harvard.edu/) to normalize the participant’s MRI structure data to the standard brain space and the cortical surface was extracted from the normalized participant brain using the polygon model with 15,002 vertices at the gray/white matter boundary. Then each vertex was assigned with a label for the corresponding ROI. Because there were exceptional vertices that did not correspond to any of the HCP-defined ROIs, which were located near the corpus callosum, these vertices were labeled by the 361st ROI (a total of 1,110 vertices were found bilaterally).

#### 2.1.6. Cortical current model

We simulated several source conditions in which the primary visual (V1) area, the inferior temporal (IT) area, the intraparietal sulcus (IPS) area, and area 8Ad were activated as representative parts of the lower visual area, the higher visual area, the lateral parietal area, and the dorsolateral prefrontal area, respectively. The duration of a single trial in the simulation was 700 ms. The period of 201–700 ms was designated as the activation phase of the target brain area. To simulate different neural activity patterns, we defined two source activation patterns. In activation pattern 1, one hemisphere was assigned a positive current of 1 nA·m, while the other was assigned a negative current of 1 nA·m. Activation pattern 2 was defined as the inverse of activation pattern 1, with the polarity of the cortical current swapped between the hemispheres.

#### 2.1.7. Sensor noise model

The noise was independently added to each MEG sensor, each time point, and each trial as shown in Eq. 1, sampled from a Gaussian distribution with a mean of 0 and standard deviation 𝜎. The value of σ was adjusted such that the resulting signal-to-noise ratio (SNR) fell within ±0.5 dB of the target SNR, which was varied from -20 dB to 10 dB with a 5 dB step, under the assumption that the signal source in V1 was activated at 1 nA·m. The SNR was calculated using the following equation:

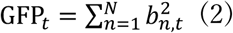

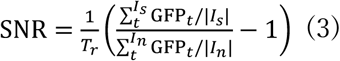

where GFP_𝑡_ represents the global field power (GFP) at time 𝑡, 𝑏_𝑛,𝑡_ denotes the MEG signal recorded from the 𝑛th sensor at time 𝑡, 𝑇_𝑟_ denotes the number of trials, and 𝐼_𝑠_ and 𝐼_𝑛_ correspond to the task and noise time windows, defined as periods during which the brain was assumed to be active and inactive, respectively. The resulting SNR was then expressed in decibels (dB) using the conversion formula 10 log_10_(SNR). The SNR was calculated after a low-pass filter with a 100-Hz cutoff frequency was applied to MEG signals with the noise. To compute the SNR, the noise and the task time window was defined as 1-200 ms and 201-700 ms, respectively. We used the sensor noise with the same magnitude calculated when V1 was active to keep the noise magnitude constant for the specified SNR levels for all source conditions. One simulation dataset consisted of five runs, each containing ten trials per activation pattern. To assess the variability of the performance of the MEG source estimation model, we made 20 datasets for each SNR by repeatedly creating the simulation dataset with different random noise patterns.

### 2.2. Real data analysis

We also analyzed real data and evaluated the performance with the following procedures.

#### 2.2.1. Participants

Three participants participated in the MRI experiment to acquire anatomical images of their brains for the extraction of the cortical surface model and the MEG experiment to acquire brain activity signals. The participants gave written informed consent before the experiments. Experimental procedure was approved by the institutional review board of the University of Electro-Communications, ATR Brain Activity Imaging Center, and Ethical Review Board of Osaka University Hospital. The experiment was conducted under corrected vision with MRI- and MEG-compatible experimental glasses, respectively.

#### 2.2.2. MRI experiment and cortical surface extraction

MRI experiments to acquire anatomical images of the participant brains and the extraction of the cortical surface model were performed using the same protocol as explained in Simulation section.

#### 2.2.3. MEG signal acquisition

MEG signals were acquired while the participants observed visual stimuli presented inside the MEG scanner. The same MEG scanner system was used as explained in Simulation section. Measurement of participants’ head position and its registration to MRI anatomical images were performed by the same procedure as for the simulation explained above.

#### 2.2.4. Visual stimulus presentation

A wedge with black-and-white checkerboard patterns was used as the stimulus in the MEG half-field visual stimulation experiment (Fig. 1b). Thus, visual stimulus conditions were twofold: stimulation of the left visual field and the right visual field. Each condition is expected to activate contralateral hemisphere in V1. The wedges had a diameter of 18 × 18 degree in visual angle and was presented on a gray background subtending 25 × 25 degrees. The stimulus sequence began with the presentation of a fixation point consisting of concentric circles at the center of the screen for 3000 ms. Then, the checkerboard wedge was presented on either the left or the right side of the visual field for 500 ms, followed by the gray background with the fixation point for 1000 ms. This wedge-background presentation was repeated three times. The color of the fixation point changed to red 500-1200 ms before the wedge presentation and returned to its original black-and-white color immediately after the last background presentation. This sequence constituted one stimulus presentation cycle. In every set of two consecutive cycles (six trials in total), the number of left and right wedge presentations was balanced such that each side was presented three times. One continuous run consisted of 28 repetitions of the cycle, and each participant completed a total of 10 runs. These visual stimuli were made and presented by Psychtoolbox-3 (http://psychtoolbox.org/).

**Fig. 1.**
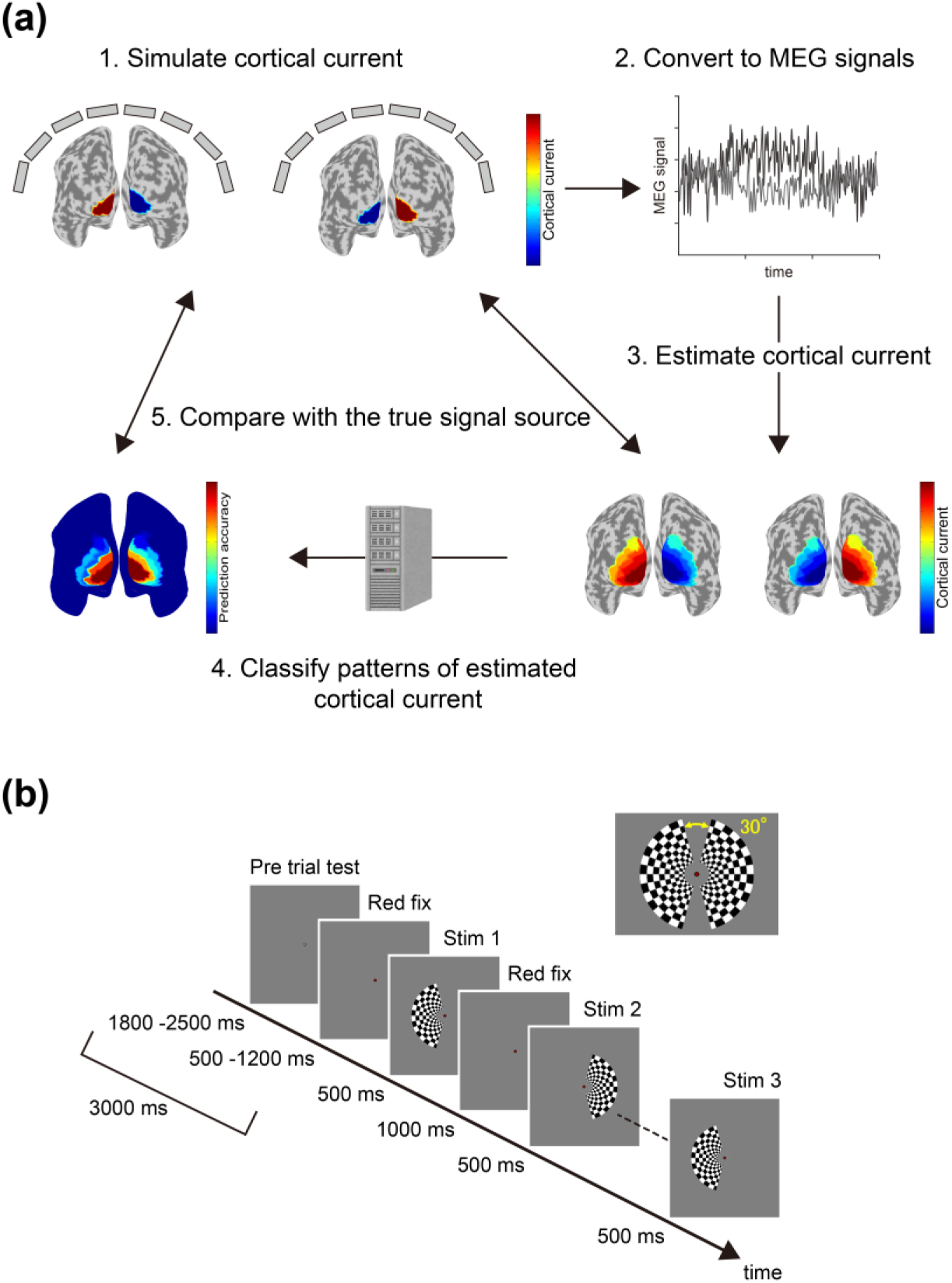
Schematic view for evaluation processes of accuracy of the cortical current estimated by MEG source estimation and stimulus prediction by decoding. (a) The cortical current was set in a specified cortical area (an example of V1 is shown) and converted to MEG signals Then, cortical currents were estimated from the MEG signals using a source estimation model. The estimated current was decoded to examine whether they contained information about the experimental conditions. Finally, the estimated current and decoding results were compared with the simulated source cortical current. (b) Experimental design of half-field visual stimuli. The wedge visual stimulus and fixation point were presented sequentially three times, comprising one cycle of stimulus presentation. One run consisted of 28 cycles, and all participants performed 10 runs.

#### 2.2.5. Preprocessing of the real data

As a preprocessing step, we first applied time-shifted principal component analysis (Cheveigné and Simon, 2007) to suppress environmental noise and a 100-Hz low-pass filter to the MEG signal. Then, the onset of the visual stimulus (checkerboard wedge) was set as 0 ms, and the period from -200 ms to 700 ms was defined as a single trial. Trials in which MEG signals exceeded ±3 pT were considered to contain movement-related or environmental noise and were excluded from further analysis. The last presentation cycle of each run was excluded from the analysis because a message indicating to stop or continue the experiment was displayed in the upper left corner of the screen. In addition, the preceding cycle was also excluded to balance the number of presentations across visual stimulus conditions.

### 2.3. MEG source estimation

MEG source estimation is to estimate 𝐣 given 𝐛 and 𝐆 in Eq. (1). However, since the number of vertices is much larger than the number of MEG sensors, a unique solution cannot be obtained (ill-posed problem). To resolve this issue, various models have been proposed. In this paper, we used the following five representative models and compared the performance in terms of source estimation accuracy and the degree suppression of information spreading.

#### 2.3.1. L2-norm regularization

A typical example to relax ill-posedness of the MEG source estimation is L2-norm regularization, or MNE (Hämäläinen, M S and Ilmoniemi, R J, 1994). This model defines the objective function as

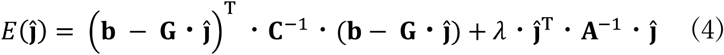

where 𝐂 is the ℝ^𝑁×𝑁^ noise covariance matrix, 𝐀 is the ℝ^𝑀×𝑀^ source covariance matrix, 𝜆 is a regularization parameter, and T indicates the transpose. The first term on the right-hand side of Eq. (2) denotes the estimation error weighted by the noise covariance matrix. The second term indicates the regularization term of the L2 norm weighted by the source covariance matrix. In this study, 𝐂 was calculated using the MEG signals from −200 ms to −1 ms in each trial. 𝐀 was set as an identity matrix. We used the MNE implementation of the Source Localization Collections (SOLCO; https://vbmeg.atr.jp/v22/toolbox/), which supports several source estimation methods used in literatures and provides files and function formats compatible with VBMEG. Among the several criteria implemented in SOLCO, we used the Akaike Bayesian information criterion to optimize the regularization parameter 𝜆 in our study.

#### 2.3.2. L1-norm regularization

L2-norm regularization tends to estimate non-zero current spread over the entire cortical surface. To suppress such broad source leakage, a sparse model is frequently used for MEG source estimation. A representative example is L1-norm regularization, also known as MCE or the least absolute shrinkage selection operator (Lasso)(Tibshirani, 1996). This model defines the objective function as

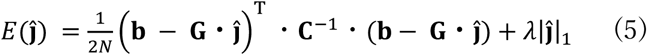

where 𝜆 is a regularization parameter and |𝑥|_𝑝_ represents the Lp-norm of a vector 𝑥. In Eq. (3), if 𝜆 is sufficiently large, the estimated current 𝐣^ is typically sparse. In this study, we tested the regularization parameter 𝜆 , ranging from 1.0 × 10^−13^ to 1.0 × 10^−1^ in exponential steps. We used the L1-norm minimization as implemented in the scikit-learn software package (http://scikit-learn.org/stable/).

#### 2.3.3. Grouped least absolute shrinkage and selection operator (gLasso)

The gLasso (Yuan and Lin, 2006) is an extension of Lasso, using the L2 norm for each group as a regularization term. The objective function is defined as

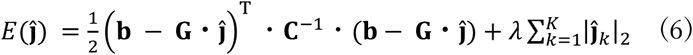

where 𝐾 is the number of groups and 𝐣_𝑘_ is the cortical current in the 𝑘th group. Lasso promotes sparsity for each cortical current whereas gLasso promotes sparsity for the group level. In this study, we tested the regularization parameter 𝜆, ranging from 1.0 × 10^−13^ to 1.0 × 10^−1^ in exponential steps. We used 180 functional regions per hemisphere, resulting in a total of 360 regions across both hemispheres (Glasser et al., 2016), along with one additional region that was not assigned to any of these 360 ROIs (in total, 361 regions for the whole brain). We used gLasso as implemented in the published code (https://github.com/AnchorBlues/GroupLasso).

#### 2.3.4. Automatic relevance determination (ARD)

ARD is a Bayesian model (Bishop, 2006; MacKay, 1992) that yields a sparse cortical current. This model introduces prior information for 𝐣 as a prior probability following a zero-mean Gaussian distribution:

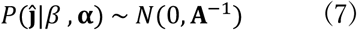

where 𝛽 represents the sensor noise variance and 𝐀 is the ℝ^𝑀×𝑀^ source covariance matrix, whose diagonal elements are 𝛼_𝑖_ (𝑖 = 1,2, … 𝑀) corresponding to the inverse of prior variance for each vertex. Note that the forward model is the same as Eq. (1). The posterior distribution of the estimated current 𝐣^ is then described as

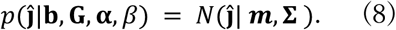

The mean 𝒎 and covariance 𝚺 are given by

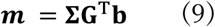

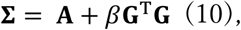

respectively. The estimated current 𝐣^ is obtained by calculating the expectation of this posterior distribution. ARD optimizes 𝛼_𝑖_ and 𝛽 through the following update rules:

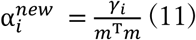

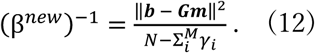

𝛾_𝑖_ is defined by

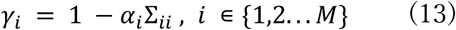

in which Σ_𝑖𝑖_ is the 𝑖-th diagonal component of Σ. Theoretically, when these update rules are repeated and 𝛼_𝑖_ becomes infinite, the estimated current ĵ_𝑖_ is converged zero (Faul and Tipping, 2001). However, in reality, when 𝛼_𝑖_ becomes large, an error occurs before 𝛼_𝑖_ reaches infinity because of the numerical limits of the software. To resolve this problem, we set the threshold of 𝛼_𝑖_ as 1.0 × 10^26^ and we considered 𝛼_𝑖_ reaching to infinity if it exceeded the threshold. If an error still occurred with this threshold value, it was progressively decreased by a factor of 10 until no error occurred.

#### 2.3.5. Functionally-structured automatic relevance determination (fsARD)

We developed a Bayesian model that incorporated functional structure information into ARD. We call this model functionally-structured automatic relevance determination (fsARD). While ARD assigns an inverse variance parameter for each vertex independently, fsARD sets a single inverse variance parameter for all vertices in each group, and updates the parameter for each group.

fsARD introduces prior information for 𝐣^ as a prior probability, 𝑃(𝐣^|𝛽 , 𝛂) ∼ 𝑁(0, 𝐀^−1^), as in Eq.(5). The source covariance matrix 𝐀 is a diagonal matrix similar to ARD, but its diagonal elements representing the inverse of the current variance are shared within the group as

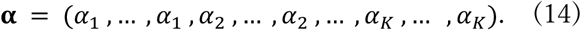

where 𝐾 represents the number of groups. The posterior distribution of the estimated current 𝐣^ is the same as that described in Eq. (6). The mean and covariance are the same as those described in Eqs. (7) and (8), respectively, but the update rules are as follows:

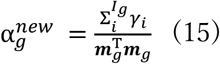

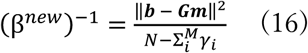

where 𝐼_𝑔_ represents the index of the vertex belonging to the 𝑔-th group. 𝛾_𝑖_ is computed by Eq. (11). Similarly to the case of ARD, we set the threshold of 𝛼_𝑘_ as 1.0 × 10^26^ and we considered 𝛼_𝑘_ reaching to infinity if it exceeded the threshold. If an error still occurred with this threshold value, it was progressively decreased in the same way as in the case of ARD. The grouping was defined based on the 361 ROIs from the HCP dataset, which are the same as the group definitions in gLasso.

### 2.4. Evaluation of source estimation accuracy and the spatial extent of information spreading

#### 2.4.1. Area under the precision–recall curve

We used the area under the precision–recall curve (APR) (Davis and Goadrich, 2006) to evaluate the accuracy of the source estimation. The area under the receiver operating characteristic curve (AUC) is commonly used for evaluation of prediction performance (Hanley and McNeil, 1982), but the previous study (Saito and Rehmsmeier, 2015) showed that APR is more suitable than AUC when the number of positive and negative values differs significantly. In this study, since the number of active vertices and inactive vertices differed significantly (imbalanced datasets), we adopted APR as an appropriate metric for the performance evaluation. Before calculating the APR, the estimated current from each model was averaged across all trials for each activation pattern. The absolute values of the averaged estimated current were normalized by the maximum of its absolute value. Using the normalized estimated current, we evaluated the presence or absence of the estimated current at each vertex while varying the threshold: if the normalized estimated current was greater than the threshold, the current was considered present at the vertex (positive case); otherwise, it was considered absent (negative case). Precision and recall were then computed at each threshold by comparing the results with the true source current distribution. Finally, we obtained a curve based on the precision and recall (precision-recall curve) and then calculated the area under the curve.

#### 2.4.2. Searchlight decoding

We conducted searchlight decoding (Kriegeskorte et al., 2006) to investigate the spatial extent of information spreading produced by MEG source estimation. Searchlight decoding identifies brain areas in which the estimated current is informative to predict experimental conditions. In our case, experimental conditions correspond to two activation patterns in the simulation data and two visual stimulus conditions in the real data. In this approach, a small target patch is defined on the cortical surface, and a decoder is trained using the estimated current within the target patch and then tested whether the experimental conditions can be predicted. This process is repeated across the entire cortex until all vertices are selected as the center of the target patch. In this study, we set each target patch consisting of 25 vertices closest to each vertex. Support vector machine (SVM) (Cortes and Vapnik, 1995) was used as decoders. This approach allowed us to compute the prediction performance at all vertices on the cortical surface and investigate the spatial extent of information spreading.

#### 2.4.3. Matthews correlation coefficient

We used Matthews correlation coefficient (MCC) (Baldi et al., 2000; Gorodkin, 2004; Matthews, 1975) to evaluate the suppression of information spreading. The F1 measure (Dice, 1945; T. Sørensen, 1948; Van Rijsbergen, 1974, 1977), also known as Dice coefficient, is widely used for evaluation of binary classification, but the previous study (Chicco and Jurman, 2020) showed that MCC is more appropriate than the F1 measure for imbalanced datasets.

In this study, since the number of predictive vertices and unpredictive vertices differed significantly, we adopted MCC as an appropriate metric for the performance evaluation. When calculating the MCC for the searchlight decoding results, we set the threshold based on the statistical significance level (p < 0.05) of the binomial test to determine whether information about the experimental conditions was present or absent at each vertex: if the prediction accuracy exceeded the threshold, the vertex was considered to contain information about the experimental conditions (positive case); otherwise, it was considered uninformative (negative case). The confusion matrix was then made by comparing the results with the true source current distribution, under the assumption that the source area contained information about the experimental conditions. Using this confusion matrix, the MCC was calculated in the same as the previous study.

#### 2.4.4. Simultaneous evaluation of source estimation accuracy and information

spreading Source estimation accuracy and suppression of information spreading were quantified using APR and MCC, respectively. To evaluate both metrics simultaneously, we plotted these values for each model as a single point in two-dimensional plane, where the horizontal axis represents APR and the vertical axis represents MCC. This plot served as an integrated visualization to assess how well each model satisfied both criteria. However, some models may exhibit imbalanced performance, achieving a high score in one metric while performing poorly in the other. For example, a previous study (Sato et al., 2018) showed that sparse models such as MCE appeared to suppress information spreading, which was actually due to poor source estimation accuracy. In such cases, the evaluation of models may vary depending on whether more emphasis is placed on source estimation accuracy or on the suppression of information spreading, making direct comparison between models difficult. To comprehensively evaluate both aspects, we computed the Euclidean distance from the origin to the point that represents each model’s APR and MCC scores. This distance served as a metric that reflects how closely each model approaches the ideal performance, where both APR and MCC are maximized.

The Euclidean distance metric was also used to determine the hyperparameters for MCE and gLasso. For both models, we explored a range of values for the regularization parameter λ. The results for all tested 𝜆 are shown in Supplementary Materials (Fig. S1 and S2). Based on the combined performance in terms of source estimation accuracy and suppression of information spreading, we selected the results with 𝜆 = 1.0 × 10^−9^ for MCE and that with 𝜆 = 1.0 × 10^−10^ for gLasso as the representative result shown in the main text. The matrix 𝐂 was set to the identity matrix in all cases.

## 3. Results

### 3.1. Simulation data analysis

#### 3.1.1. Mapping of estimated current and searchlight decoding accuracy

First, we examined the maps of the normalized estimated current and searchlight decoding accuracy to qualitatively evaluate the source estimation accuracy and the spatial extent of information spreading.

Figure 2 shows the normalized estimated current maps by each model for activation pattern 1 under three representative SNR levels. When the signal source was located in V1 (Fig. 2, upper left), MNE with squared regularization produced spatially spread solutions, resulting in nonzero estimated currents across the entire cortical surface. This indicates widespread leakage of the cortical current beyond the source area. In contrast, the estimated currents obtained with MCE and ARD, both of which promote sparse solutions, showed sparse cortical current, with many cortical vertices exhibiting zero values, leading to a narrow spatial extent of source leakage. gLasso, which incorporated functional structure information as regularization terms, resulted in a narrower spatial extent of source leakage compared to MNE for higher SNR levels. However, gLasso exhibited nonzero estimated currents similar to those of MNE for lower SNR levels, indicating widespread leakage of cortical current beyond the source area. fsARD, which combines Bayesian estimation with functional structure information, estimated relatively high-amplitude cortical currents localized in V1 for higher SNR levels. Although fsARD also exhibited source leakage, the leak amplitude was lower than that of MNE and gLasso. As the noise level increased, all models exhibited reduced source estimation accuracy, with either the emergence of high-amplitude estimated currents outside V1 or the expansion in the spatial extent of the estimated current, compared to lower noise levels. These results were qualitatively similar for activation pattern 2 (see Supplementary Materials (Fig. S3)).

**Fig. 2.**
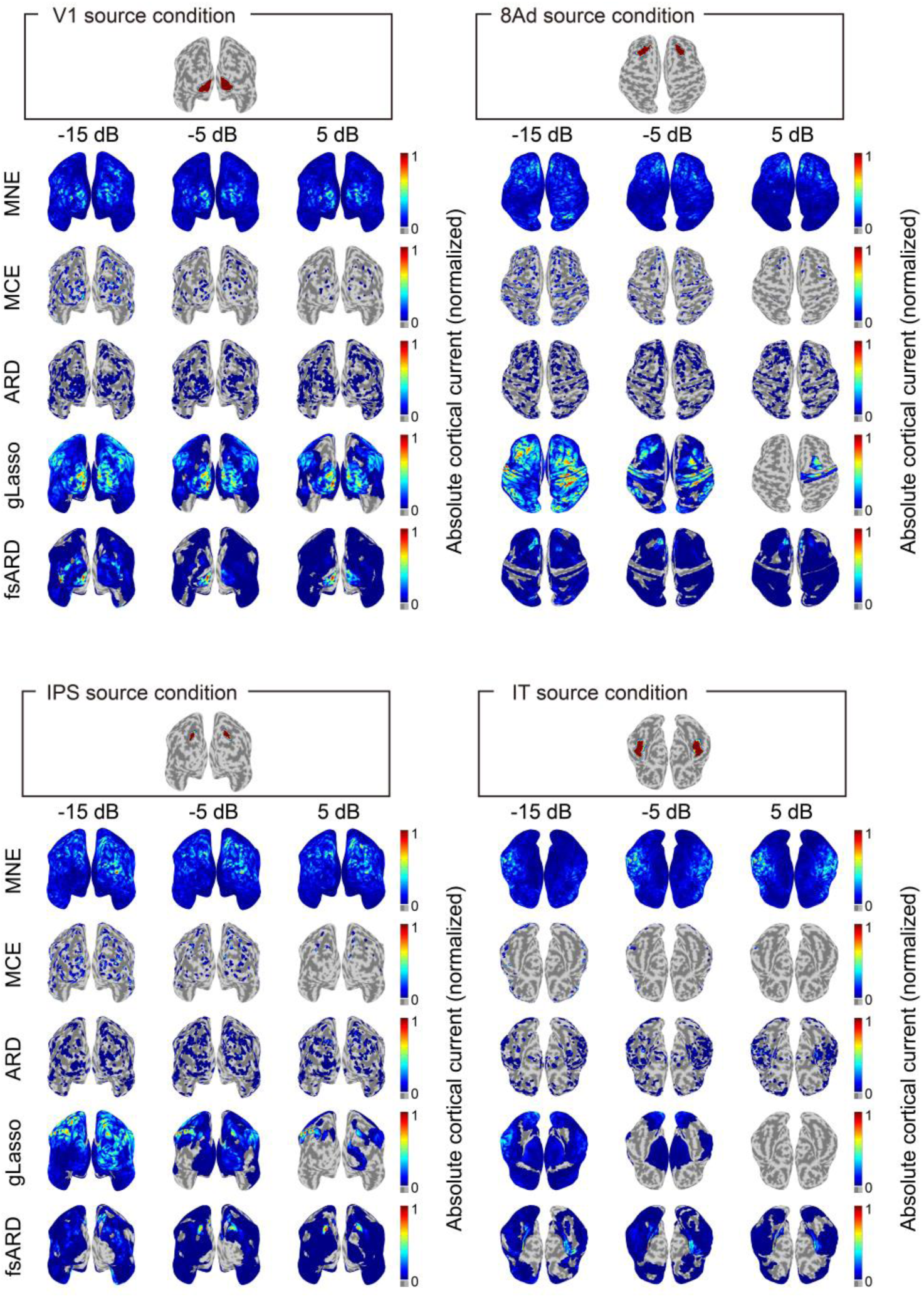
Mapping of estimated current. The results for three different SNR levels (- 15dB, -5dB, and 5dB) for activation pattern 1 are shown.

Figure 3 shows the searchlight decoding accuracy maps for each model under the same SNR levels as in Figure 2. When the signal source was located in V1 (Fig. 3, upper left), MNE produced high prediction accuracy across all vertices on the cortical surface, indicating that the estimated current at every vertex contained information about the activation patterns, as shown in Sato et al. (2018). In contrast, MCE and ARD produced low prediction accuracy across most vertices, while exhibiting high prediction accuracy in a small number of specific areas. This is because MCE and ARD estimated the cortical current to be zero in most areas, resulting in the loss of information about the activation patterns in those areas. gLasso produced high prediction accuracy across most vertices, similar to MNE, indicating that the estimated current at most vertices contained information about the activation patterns. fsARD showed high prediction accuracy in V1 while maintaining chance-level accuracy in areas outside V1, demonstrating its superior ability to suppress information spreading compared to other models. As source estimation accuracy declined at lower SNR levels, the spatial extent of information spreading also diminished, as prediction of the activation patterns became more difficult due to inaccuracies in the estimated cortical currents.

**Fig. 3.**
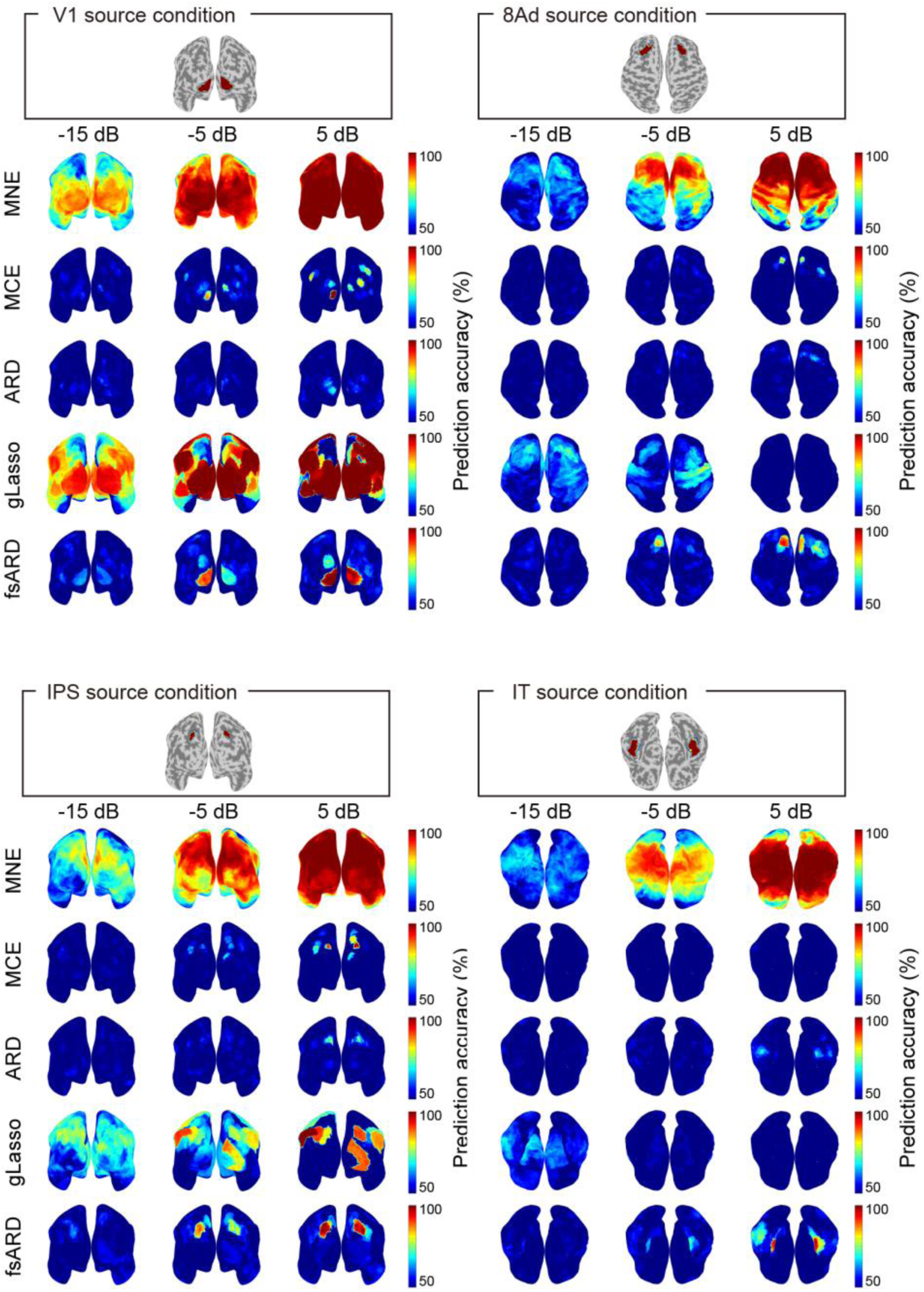
Mapping of searchlight decoding accuracy. Prediction accuracy between 50% and 100% are displayed. The results for three different SNR levels (-15dB, -5dB, and 5dB) are shown.

In the other source conditions, the spatial extent of the estimated current and information spreading varied depending on SNR levels, qualitatively similar to those observed in the V1 source condition (Fig. 2 and 3). However, the 8Ad and IT source conditions tended to reduce both source estimation accuracy and decoding accuracy, suggesting that source estimation is more difficult for these areas than for V1.

#### 3.1.2. Quantitative evaluation of source estimation accuracy and the spatial extent of information spreading

To quantitatively evaluate the source estimation accuracy and the spatial extent of information spreading of each model, we calculated the APR of the normalized estimated current and MCC of searchlight decoding results while changing the SNR level. Figure 4 shows the results for activation pattern 1, plotting the APRs of the normalized estimated current along the horizontal axis and the MCCs of searchlight decoding along the vertical axis for each model and source condition.

**Fig. 4.**
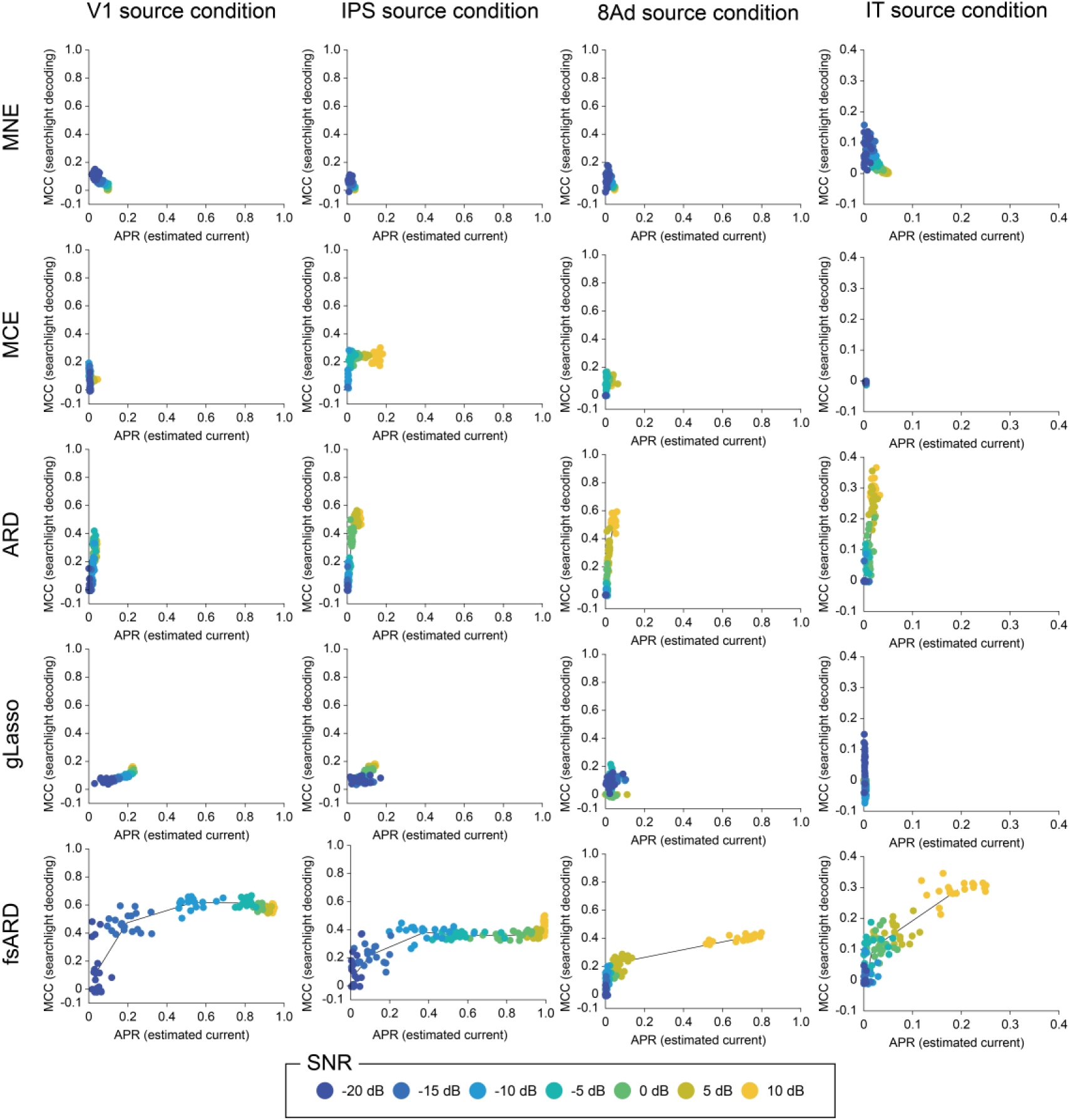
Quantitative evaluation of source estimation accuracy and suppression of information spreading. Each point represents APR of the estimated cortical current for activation pattern 1 (horizontal axis) and MCC of the searchlight decoding (vertical axis) for a given SNR level. SNR levels are indicated by the color scale shown at the bottom.

When the signal source was located in V1 (Fig. 4, left column), MNE produced an inverse relationship between the APR of the estimated current and the MCC of searchlight decoding: as the APR of the estimated current decreased with lower SNR levels, the MCC of searchlight decoding increased. This occurred because, as the noise level increased, source estimation accuracy declined and false positive predictions by decoding outside the source area (V1 in this case) decreased, leading to a pseudo-improvement in the MCC and resulting in apparent suppression of information spreading. MCE showed low APR and MCC for all tested SNRs, because it produced sparse estimation with zero cortical currents at almost all vertices. The MCC of searchlight decoding was also low because only a few vertices showed a high prediction accuracy in V1, resulting in fewer true positives. As shown in Fig. 2, ARD also exhibited sparse estimation and thus the APR of the estimated current was comparable to MCE. However, the MCC of searchlight decoding was higher than that of MCE because high predictions were observed at a larger number of vertices in V1. gLasso exhibited less source leakage compared to MNE and thus showed a higher APR of the estimated current for higher SNR levels. However, the MCC of searchlight decoding remained as low as that of MNE since it produced many false positive predictions even outside V1. In contrast to all these models, fsARD demonstrated a simultaneous increase in both the APR of estimated current and the MCC of searchlight decoding, attaining the highest values except at the lowest SNR. These results indicate that fsARD achieved the best performance regarding the simultaneous evaluation of source estimation accuracy and suppression of information spreading, in a wide range of SNR tested in this study.

Both APR and MCC exhibited similar trends with respect to SNR across all models and source conditions, although their values tended to be lower for the 8Ad and IT source conditions. These lower performance was particularly pronounced for the IT source condition, where even fsARD, which achieved the highest performance in the V1 source condition, the maximum APR and MCC reached only around 0.2 and 0.3, respectively. These results were qualitatively similar for activation pattern 2 (see Supplementary Materials (Fig. S4)).

Figure 5 shows the mean Euclidean distance from the origin to the point that represents each model’s APR and MCC scores for activation pattern 1, averaged across 20 simulated points for each SNR. A similar trend to that observed in Figure 4 was evident in Figure 5, where fsARD outperformed the other models in the V1 and IPS source conditions, whereas its performance was comparable to that of the other models in the 8Ad and IT source conditions. These results were qualitatively similar for activation pattern 2 (see Supplementary Materials (Fig. S5)).

**Fig. 5.**
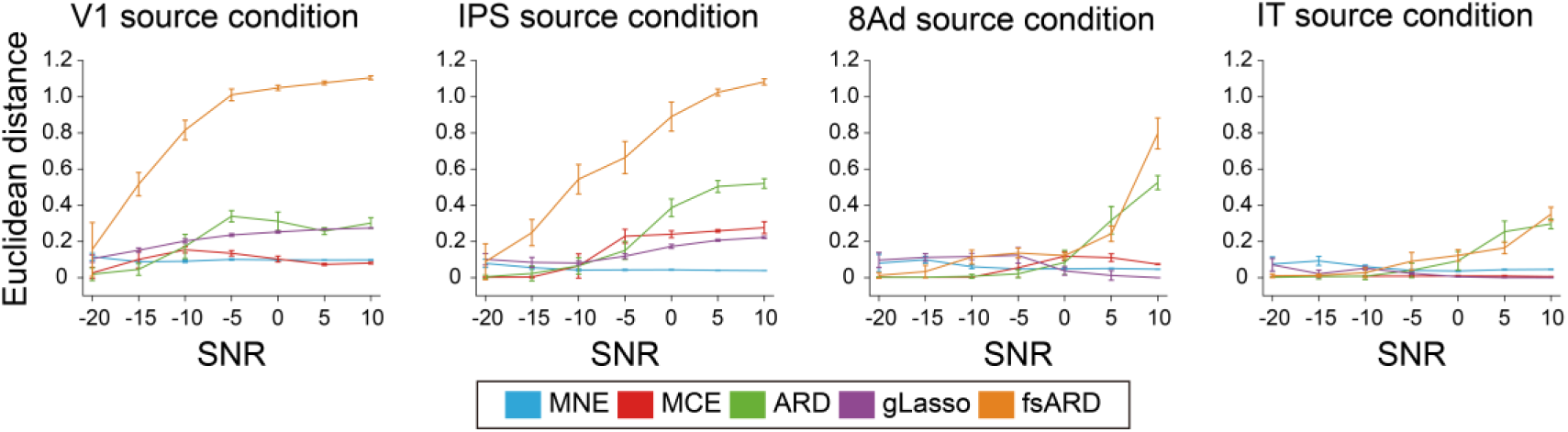
Euclidean distance from the origin to the point that represents each model’s APR and MCC scores across SNR levels. Each line shows the mean of the Euclidean distance from the origin (vertical axis) in the APR-MCC space under activation pattern 1, computed across 20 dataset and plotted against the SNR level (horizontal axis). Error bars indicate the standard deviation. Each model is indicated by the color scale shown at the bottom.

### 3.2. Real data analysis

To evaluate model performance with the real data, we conducted MEG experiments using visual stimuli and applied the MEG source estimation models to the measured data. In this experiment, V1 in the left hemisphere was assumed to be activated when the visual stimulus was presented in the participant’s right visual field, and vice versa. We calculated the GFP of the MEG signal during the stimulus presentation period and estimated the cortical current at the time point corresponding to the GFP peak. This peak occurred at 84 ms, 94 ms, and 95 ms after stimulus onset for each participant, respectively. These timings correspond to feedforward neural activity evoked in V1 according to the consistency of the latency of the human early visual area (Yoshor et al., 2007). Based on this correspondence, we assumed that the V1 was the activity source for each visual stimulus condition, and used this assumption as the basis for the subsequent evaluation. As in the simulation analysis, the estimated current from each model was averaged across all trials. The averaged estimated current was then normalized by its maximum value. Using this normalized estimated current, we calculated the APR for each visual stimulus condition (i.e., left and right wedge presentations).

Figure 6 shows estimated current maps obtained by the tested models. Results were qualitatively similar to those of simulation. MNE produced spatially spread estimation, resulting in nonzero estimated current across almost all vertices beyond the true source area. In contrast, MCE and ARD produced sparse estimation, resulting in the estimated current being zero at most of the vertices both inside and outside V1. gLasso showed spatially spread current estimation as MNE but the spatial extent of source leakage beyond V1 was narrower. The magnitude of the estimated current in V1 was greater than that of the other models, but substantial current was also estimated outside V1, even more so than in the case of MNE. fsARD was able to estimate current in V1. Source leakage was also observed across widespread areas, but its spatial extent was smaller than that of MNE and gLasso.

**Fig. 6.**
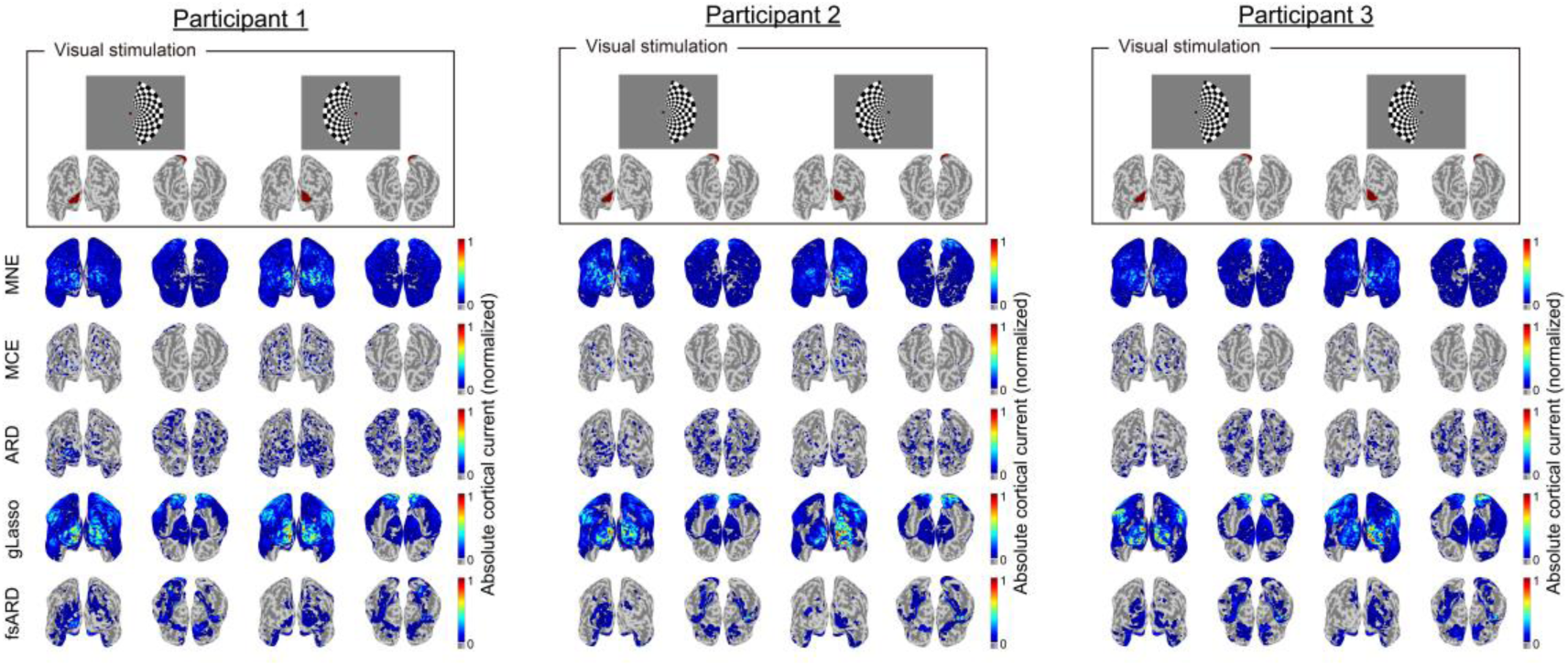
Mapping of estimated current in real data analysis. The normalized estimated current maps are shown for each model, each participant, and each visual stimulus condition.

Next, we conducted searchlight decoding to investigate the spatial extent of information spreading. The estimated current by each model was used as input to classifiers to discriminate between the two visual stimulus conditions (left or right visual field stimulation). When visual stimuli are presented in either the left or right visual field, the V1 hemisphere on the opposite side of the stimulated hemifield is activated. Therefore, both bilateral V1 are expected informative to predict the visual stimulus conditions when searchlight decoding is performed.

Figure 7 shows the maps of the decoding accuracy for each participant, indicating qualitatively similar results to the simulation. MNE exhibited high prediction accuracy across almost all vertices. In contrast, MCE and ARD showed low prediction accuracy in the broad cortical areas except a small number of vertices where the estimated cortical current was nonzero. gLasso exhibited high prediction accuracy across most vertices similar to MNE. fsARD showed higher decoding accuracy in the V1 and its surroundings, but it was near the chance level in most areas outside V1.

**Fig. 7.**
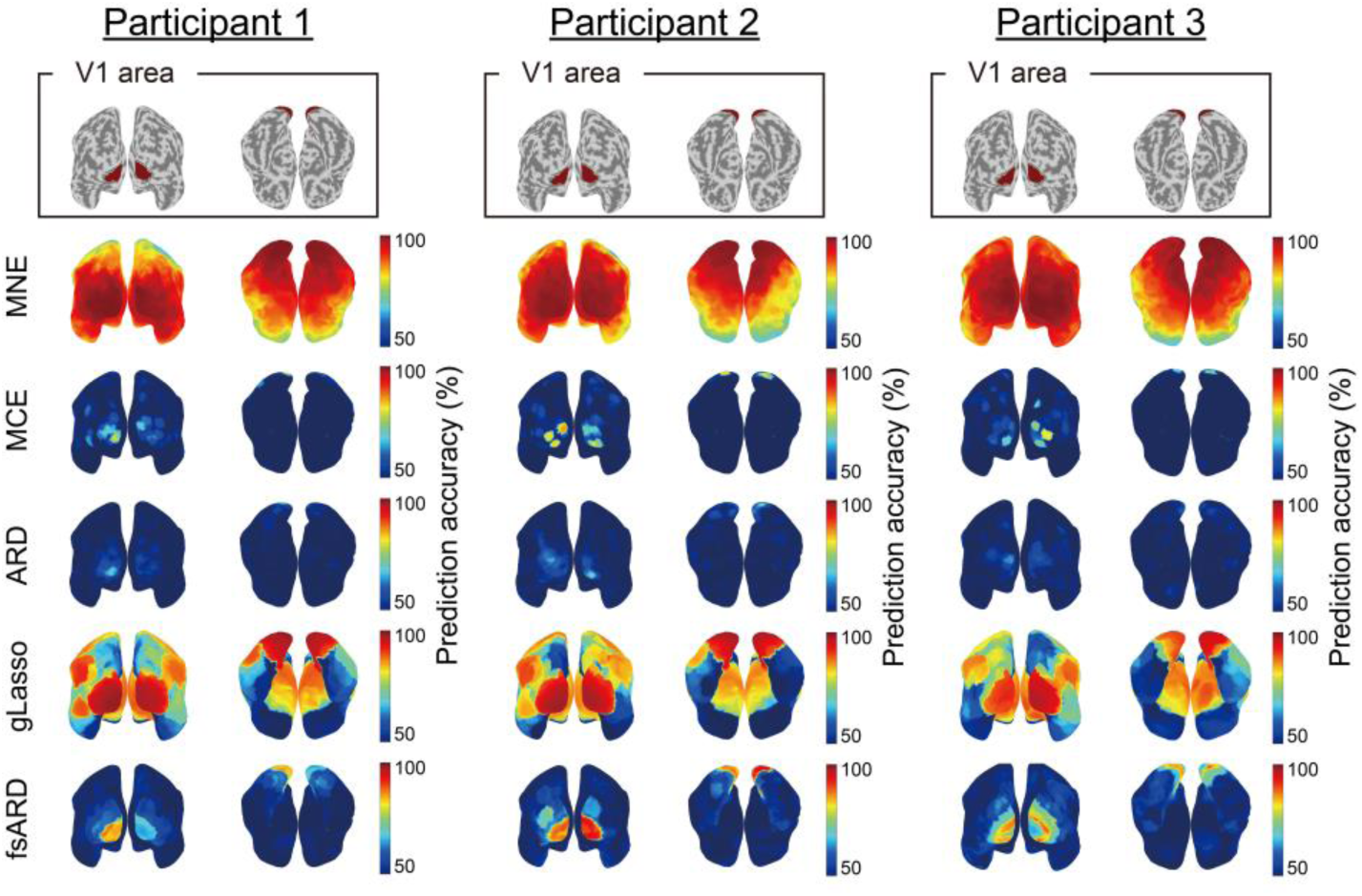
Mapping of searchlight decoding accuracy in real data analysis. Prediction accuracy between 50% and 100% are displayed.

To quantitatively evaluate these results, we calculated APR for the estimated current and MCC searchlight decoding. To calculate APR, we used the normalized estimated current. Then, as in the simulation, we varied a threshold to determine presence or absence of the cortical current at each vertex in the range from 0 to 1. In this study, we assumed that V1 hemisphere contralateral to the presented visual stimulus was activated while the other areas was not, as the visual stimulus was the simple checkerboard wedge and V1 is expected to be most active in the cortical areas. We hypothesized this current distribution as a “true source current,” and compared it with the results of the estimated current to compute precision and recall at each threshold. Finally, we obtained a precision-recall curve and calculated the area under the curve. MCC was also similarly calculated as in the simulation. The confusion matrix was then made by comparing the results with the true informative cortical area. Here we defined it as bilateral V1 because our visual stimulus is expected to activate either left or right V1 hemisphere and thus the bilateral V1 activation status should be informative to predict the visual stimulus condition.

Figure 8 shows the simultaneous evaluation of source estimation accuracy and information spreading by plotting APR and MCC scores (panel a) and the Euclidean distance from the origin to the point that represents each model’s APR and MCC scores (panel b). These results indicate that fsARD achieved the most balanced performance among the tested models, exhibiting moderate source estimation accuracy and the highest decoding specificity. As a result, it showed the longest Euclidean distance from the origin to the point that represents fsARD’s APR and MCC scores, reflecting its strong overall performance across both evaluation metrics.

**Fig. 8.**
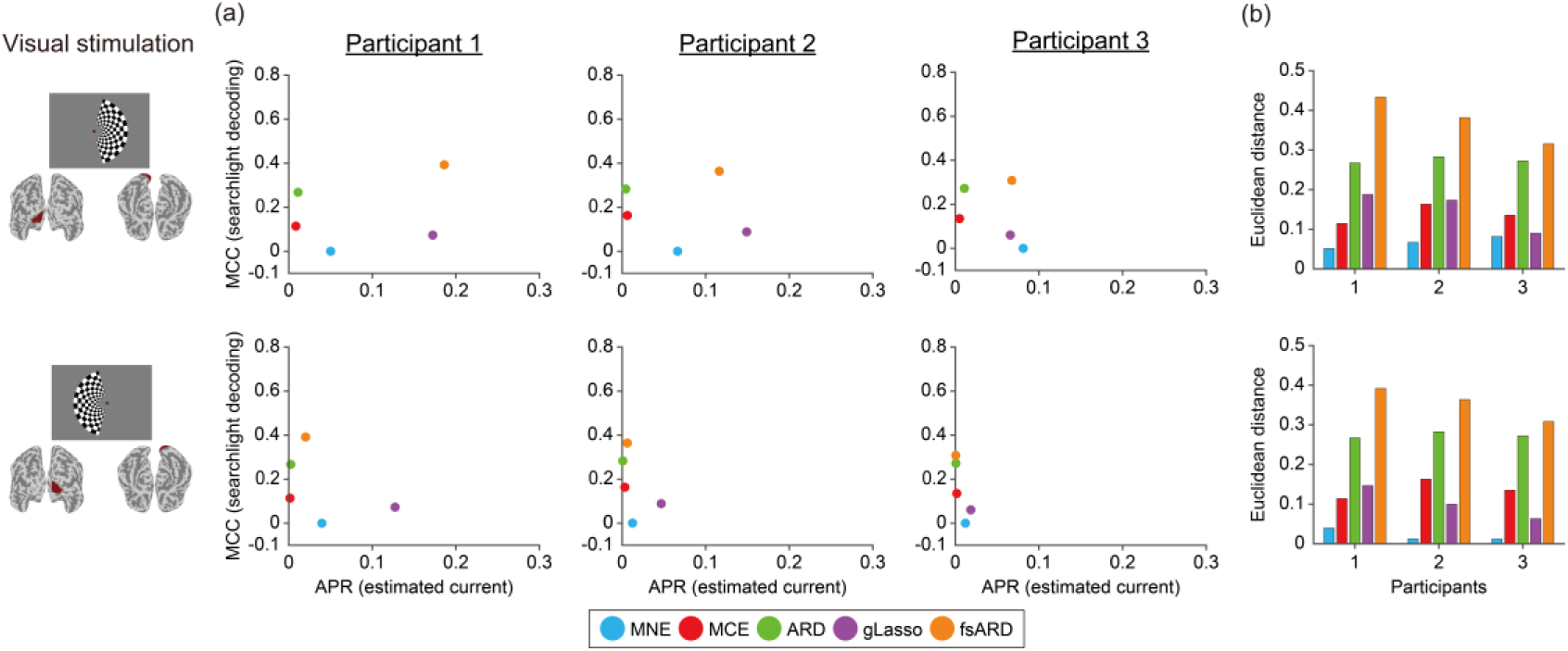
Quantitative evaluation of source estimation accuracy and suppression of information spreading in real data analysis. (a) Each point represents APR of the normalized estimated cortical current (horizontal axis) and MCC of the searchlight decoding (vertical axis) for each model. (b) Euclidean distance from the origin to the point that represents each model’s APR and MCC scores. Each bar represents the Euclidean distance (vertical axis) and participants (horizontal axis) for each model.

## 4. Discussion

In this study, we proposed a functionally-structured Bayesian model to suppress information spreading while maintaining high source estimation accuracy in MEG source estimation. Unlike previous models (Hämäläinen, M S and Ilmoniemi, R J, 1994; MacKay, 1992; Sato et al., 2004; Tibshirani, 1996), which treated each vertex independently and disregarded the functional organization of the brain, the proposed model enables estimations that reflect the similarity of activity in the same functionally-defined area. This feature is achieved by incorporating functional structure information derived from fMRI data (Glasser et al., 2016) and by estimating the prior variance parameter on an area-by-area basis. Results from both simulations and real MEG data demonstrated that the proposed model outperforms the existing models in both source estimation accuracy and suppression of information spreading.

The results of MEG source estimation show that the proposed model can estimate cortical current with a reasonable degree of sparseness by incorporating functional structure information of the brain. Among the models used in this study, MCE and ARD also produced sparse estimations; however, their estimates were excessively sparse. By incorporating functional structure constraints, the proposed model enables the sharing of prior variance parameters within each functional group, which may help avoid the overly sparse estimates that are unnatural as neural activity distribution observed in the real data. Moreover, unlike MNE and gLasso, the proposed model achieved to reduce source leakage to areas where no signal source was expected (Fig. 2). This result proved that the proposed model worked as intended under the assumption that the brain activity is similar within the functionally same area. Although gLasso also incorporated the same functional structure information as fsARD, it exhibited information spreading as MNE did. This is likely because gLasso, unlike Bayesian estimation models, does not account for noise variance and might misinterpret noise as signal sources.

The results of the searchlight decoding demonstrate that the proposed model suppresses information spreading while maintaining accurate source estimation. Previous studies have reported that sparse models, such as MCE, do not cause information spreading (Sato et al., 2018). This study replicated that both MCE and ARD did not show information spreading (Fig. 3,4 and 5). However, this phenomenon is likely attributable to the excessive sparseness and inaccuracy of the cortical current estimation, from which the decoder was not able to retrieve information. The proposed model successfully mitigated this issue by applying group constraints, demonstrating accurate decoding in the true source area and its vicinity. On the other hand, accurate decoding was not possible outside the true source area, even in areas where source leakage was observed. These findings suggest that the leak current likely reflects noise components or brain activity unrelated to the activation patterns. Thus, the information is not actually spread through the cortical current estimation by the proposed model. Although gLasso also incorporates the same group constraints, it yielded the source leakage in which the leak current falsefully preserved information that could be used to discriminate the activation patterns (Fig. 2), thereby failing to suppress information spreading.

However, it is possible that the source activation patterns used in this study were more suitable for fsARD, which may have influenced the results of source estimation and searchlight decoding. In the simulation, the source cortical current was set such that all vertices within a single ROI were activated at a constant amplitude. This setting is likely advantageous for gLasso and fsARD because both assume that similar brain activity occurs in each functional area. Therefore, it might be necessary to assume a cortical current with a complex pattern for fair evaluation. To examine whether such setting is actually influential, our previous study (Koizumi et al., 2022) used a source current with a complex spatial pattern, where the current amplitude at each vertex was sampled from a Gaussian distribution, and performed source estimation and searchlight decoding using fsARD. Results showed that for higher SNR levels, fsARD successfully performed source estimation accurately and suppressed information spreading better than models without group constraints. These results suggest that the superior performance of fsARD is not merely an artifact of the specific simulation setup of the present study but holds even when the source current has the spatial complexity.

In this study, we aimed to achieve accurate source estimation and suppression of information spreading simultaneously in MEG source estimation. We consider this an important criterion because while accurate source estimation ensures the localization of brain activity, failure to suppress information spreading can result in misleading interpretations about where the neural information is represented. Our results indicate that conventional models consistently failed to maintain this balance: while some models excel in one aspect, they tend to compromise the other. For example, sparse models (MCE and ARD) suppressed information spreading, but it was largely due to poor source estimation. In another example using MNE, the degree of suppression of information spreading increased as the noise level increased, but this was due to a decrease in source estimation accuracy (Fig. 4). In contrast, the proposed model achieved balanced performance in both aspects by incorporating prior information derived from functional structure of the human brain in a Bayesian framework.

The proposed model exhibited similar trends in how APR and MCC varied with SNR levels across all source conditions. However, their values of APR and MCC tended to be lower in the 8Ad and IT areas compared to those in the V1 area. The lower performance can be partially attributed to the sensor noise level in this study. Specifically, the noise amplitude was determined so that the SNR ranged from -20 dB to 10 dB, assuming the signal source was in V1. The same sensor noise was also applied when the source was located in other areas. Consequently, the actual SNR varied across areas: approximately -25 dB to 6 dB in IPS, -28 dB to 3 dB in 8Ad, and -28 dB to 2 dB in IT. These differences in actual SNR likely contributed to the reduced model performance in other areas.

Given this vulnerability to noise, further improvements are needed to achieve high performance at lower SNR levels. In the real data, the SNRs for participants 1 to 3 were -15 dB, -12 dB, and -16 dB, respectively. At these SNRs, fsARD showed the most balanced performance among the tested models (Fig. 8). However, since this study specifically used the experimental design that activated the V1 area, the model’s performance for other ROIs has been unclear. According to the simulation results, the model’s performance decreased in other areas such as 8Ad and IT compared to V1. Therefore, for real data, the model’s performance may also decline when targeting other areas, especially those located farther from the MEG sensors than V1. Therefor, it is important to address the model’s vulnerability to noise in order to achieve high performance across a broader range of areas. Previous studies (Cai et al., 2023, 2021) have explored models to enhance noise robustness. For instance, the Champagne model (Wipf et al., 2010) improved noise robustness by learning sensor noise covariance and signal source correlations by using pre-stimulus data. This approach has been shown to improve source estimation accuracy. However, its impact on information spreading remains unclear, and thus incorporating a similar procedure into our model would be worth exploring in future studies.

Although fsARD achieved accurate source current estimation and suppression of information spreading, there remains possibilities for further improvement. First, the group definition could be further tailored to the purpose of analyses. For example, a brain atlas known as the Penfield homunculus could be used to further subdivide motor and somatosensory areas into finer functional units (Penfield and Rasmussen, 1950), such as those representing the hands, fingers, and face. Second, inter-areal interactions could be integrated into the model together with functional structure information adopted in the present study. For example, temporal and spatial correlations defined by functional connectivity between brain areas, and geodesic distances along the cortical surface could be sources of information of the inter-areal interactions. Such additional information could improve accuracy of source estimation and suppression of information spreading. With the assistance of these additional information, we might be able to analyze when and where each brain area represents perceptual information. Thus our model framework might provide new possibility to capture human neural activity patterns and their represented information at high spatio-temporal resolution.

## Data and Code Availability

The code used in this study is available at: https://github.com/k-miyazaki1224/fsARD The dataset used in this study is available from the authors upon reasonable request.

## Author Contributions

Conceptualization: KM and YM; Formal analysis: KM; Funding acquisition: KM, OY and YM; Investigation: KM, SN, NI and KA; Methodology: KM, SN, OY and YM; Software : KM, SN, NI and KA; Supervision: YM; Visualization: KM; Writing – original draft: KM; Writing – review & editing: KM, SN, NI, KA, OY and YM.

## Acknowledgments

We thank Stuart Jenkinson, PhD, from Edanz (https://jp.edanz.com/ac) for editing a draft of this manuscript.

## Supplementary Material

Supplementary material for this article is available with the online version.

