## Supplementary Materials for "Functionally-structured Bayesian model for localizing neural activity and information in magnetoencephalography signals"

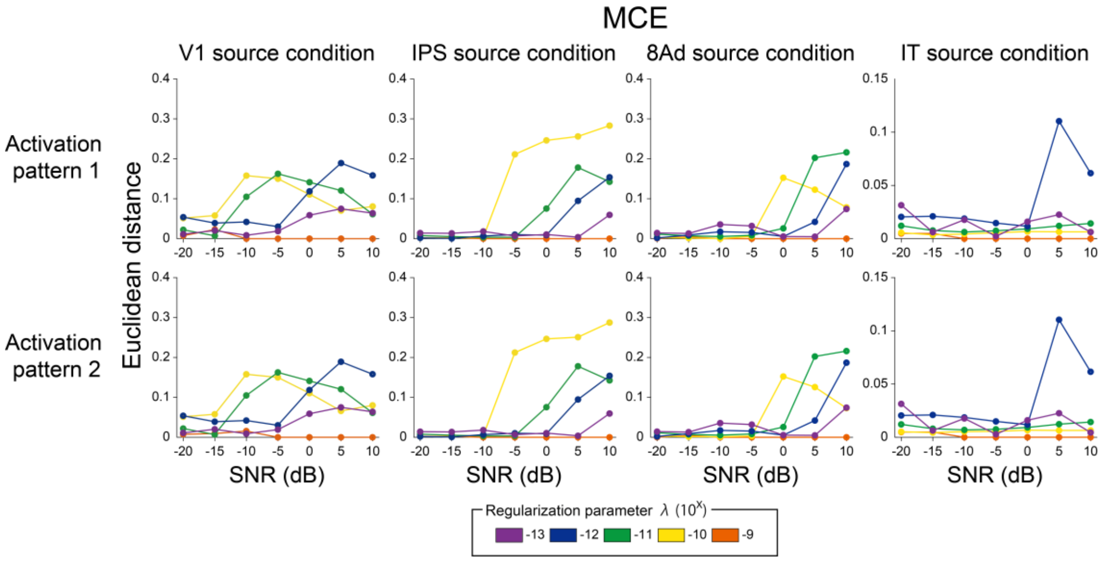


Fig. S1. **Euclidean distance from the origin to the point that represents each model's APR and MCC scores for MCE.** The regularization parameter $\lambda$ was varied in the range between $1.0 \times 10^{-13}$ and $1.0 \times 10^{-1}$ in exponential steps. However, since both APR and MCC were zero for the range from $1.0 \times 10^{-8}$ to $1.0 \times 10^{-1}$, the results for this range are not shown in the figure. Each line shows the Euclidean distance from the origin to the point that represents each model's APR and MCC scores for MCE (vertical axis), plotted against the SNR level (horizontal axis) for each source condition and source activation pattern.

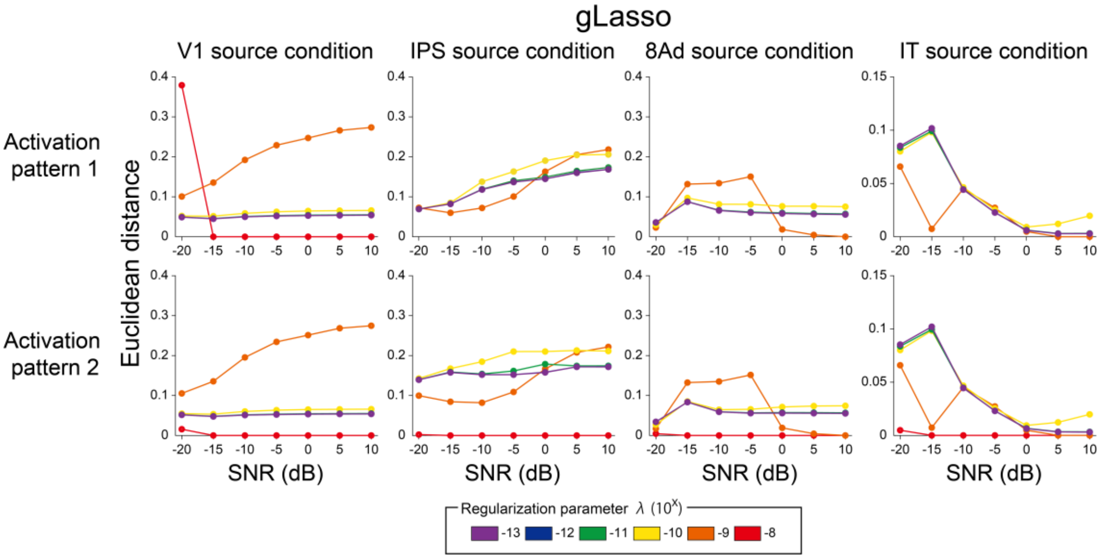


Fig. S2. **Euclidean distance from the origin to the point that represents each model's APR and MCC scores for gLasso.** The regularization parameter $\lambda$ was varied in the range between $1.0 \times 10^{-13}$ and $1.0 \times 10^{-1}$ in exponential steps. However, since both APR and MCC were zero for the range from $1.0 \times 10^{-7}$ to $1.0 \times 10^{-1}$, the results for this range are not shown in the figure. Each line shows the Euclidean distance from the origin to the point that represents each model's APR and MCC scores for gLasso (vertical axis), plotted against the SNR level (horizontal axis) for each source condition and source activation pattern.

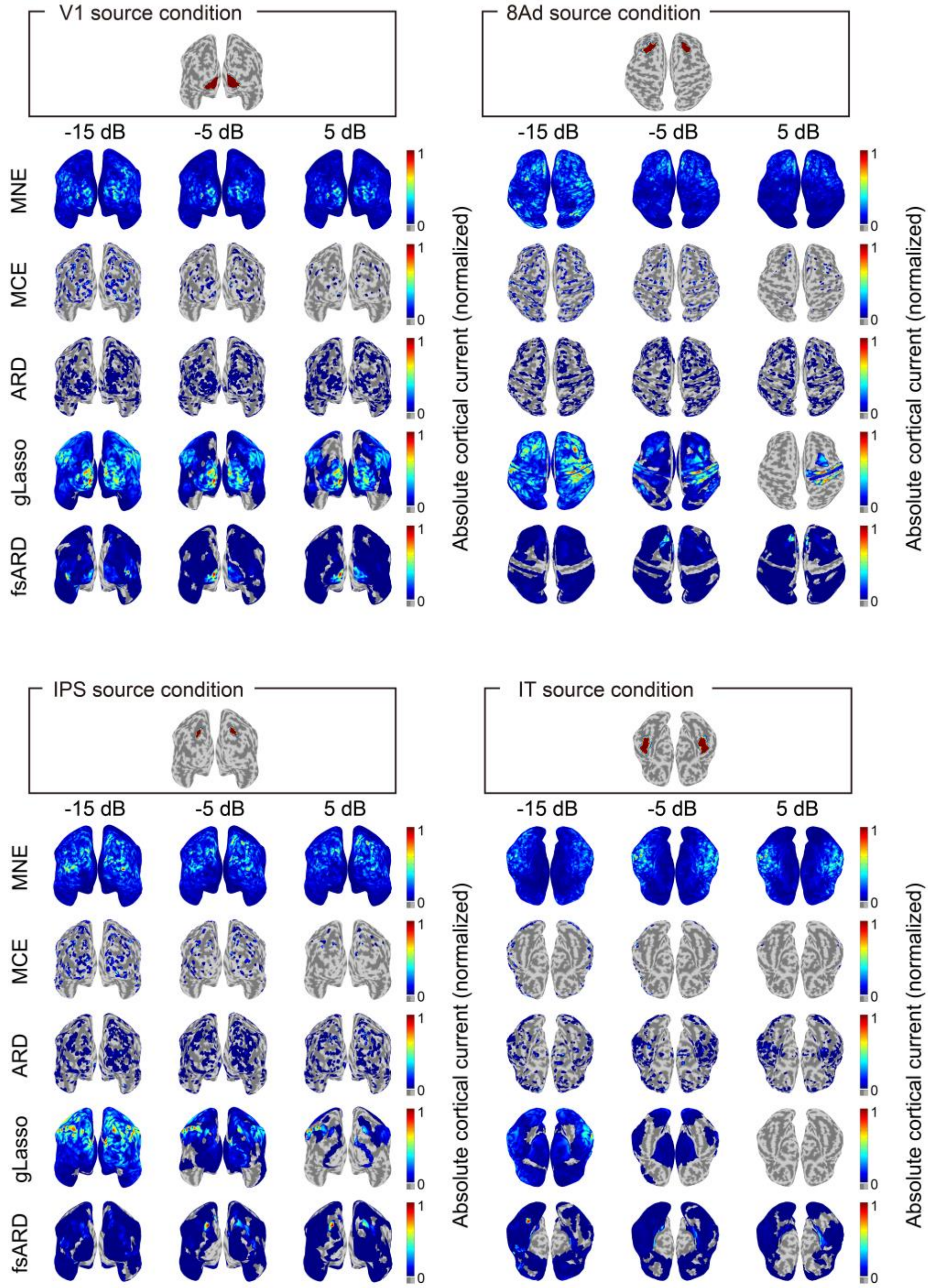


Fig. S3. **Mapping of estimated current.** The results for three different SNR levels (-15dB, -5dB, and 5dB) for activation pattern 2 are shown.

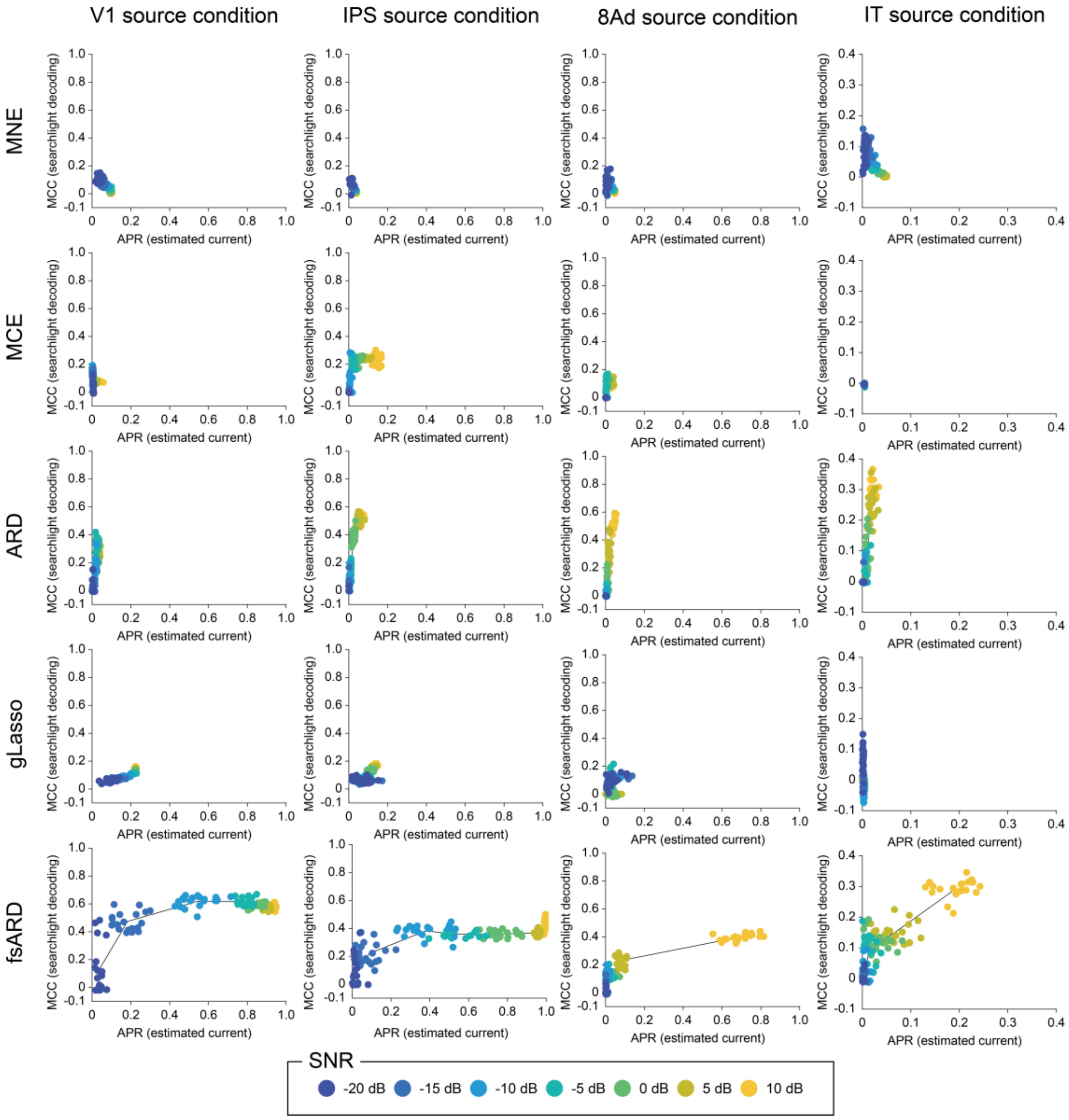


Fig. S4. **Quantitative evaluation of source estimation accuracy and suppression of information spreading.** Each point represents APR of the estimated cortical current for activation pattern 2 (horizontal axis) and MCC of the searchlight decoding (vertical axis) for a given SNR level. SNR levels are indicated by the color scale shown at the bottom.

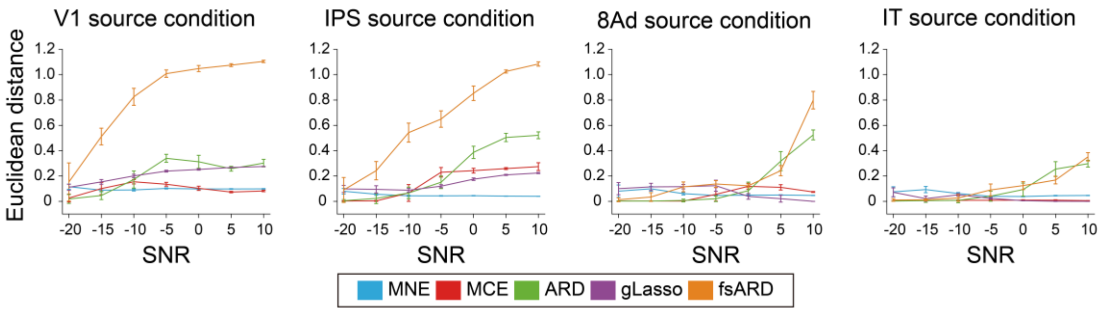


Fig. S5. **Euclidean distance from the origin to the point that represents each model's APR and MCC scores across SNR levels.** Each line shows the mean of the Euclidean distance from the origin (vertical axis) in the APR-MCC space under activation pattern 2, computed across 20 dataset and plotted against the SNR level (horizontal axis). Error bars indicate the standard deviation. Each model is indicated by the color scale shown at the bottom.
